## Supplemental Data version 2 for "Missense variant analysis in the TRPV1 ARD reveals the unexpected functional significance of a methionine"

Supplemental Figure 1

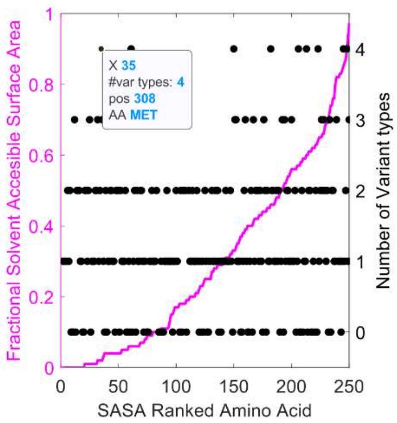


**Figure S1.** Fractional SASA score (pdb structure file: 7lp9) is plotted against SASA ranked amino acids of the ARD (residues 111-361) in rat TRPV1 (purple). Missense variant type count in gnomAD 4.1 for each position (black marker) is indicated on the right-side y-axis. SASA ranked amino acid X=35 corresponds to M308 in the rat TRPV1 sequence and as indicated on the top left box. The Matlab script for generating this plot is described in Mott et al. (2023) and available through gGitHub (*esenlab*/variant-analysis).

Supplemental Figure 2

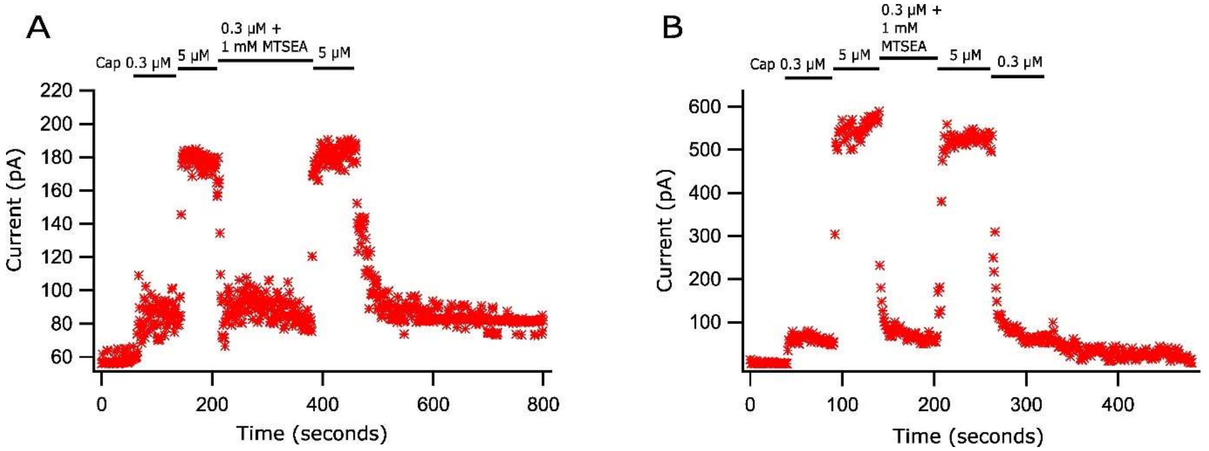


**Figure S2. Application of MTSEA to TRPV1-M308C/C157A excised patches does not affect channel activity. (A)** Macroscopic current voltage clamp experiment with application of capsaicin and MTSEA as indicated over traces. Control TRPV1-C157A with 1 mM MTSEA and co-application of 0.3 µM capsaicin. **(B)** TRPV1-C157A/M308C with 1 mM MTSEA and co-application of 0.3 µM capsaicin.

Supplemental Figure 3

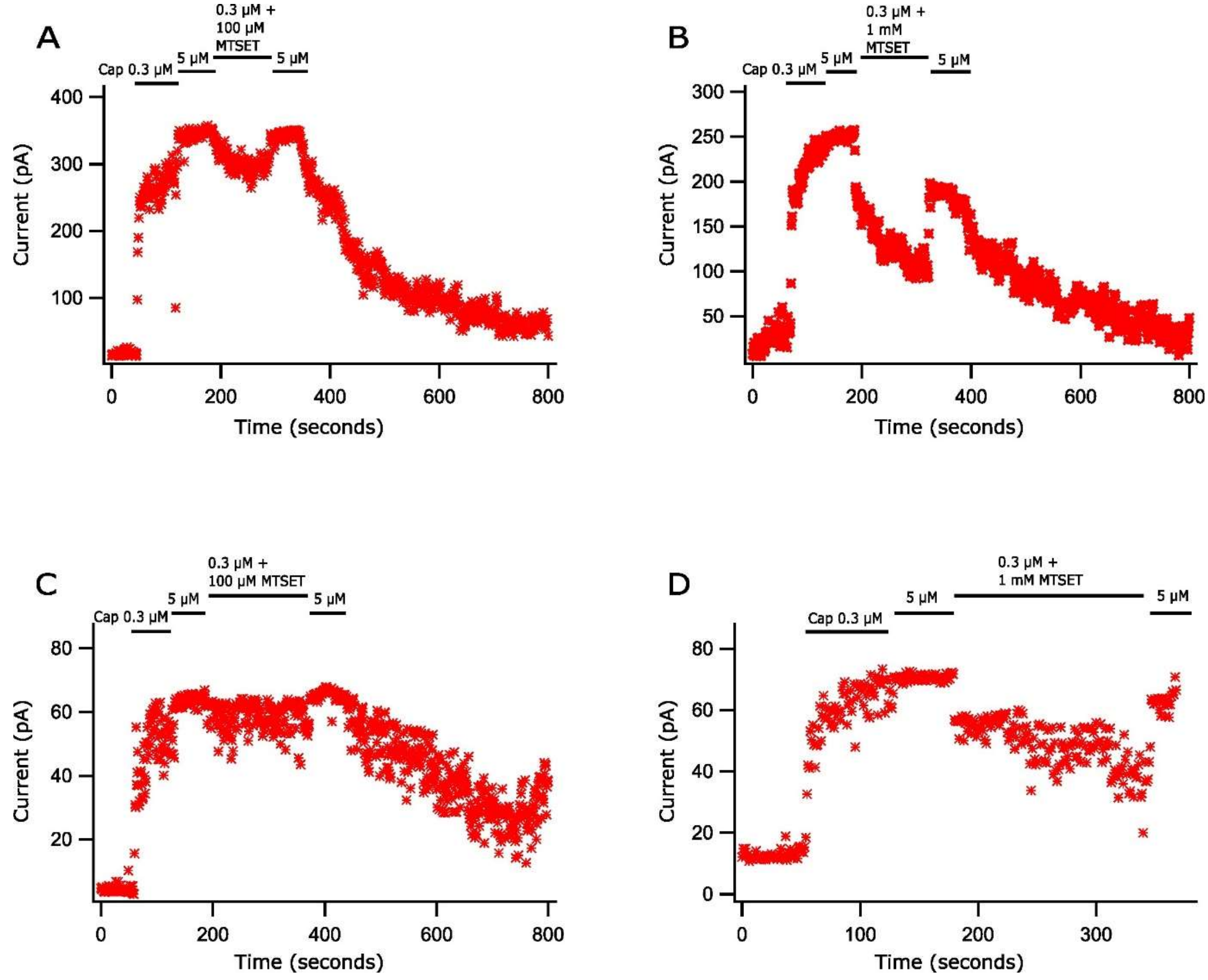


**Figure S3. Application of MTSET to TRPV1-M308C/C157A-expressing excised patches does not affect channel activity. (A)** Macroscopic current voltage clamp experiment with application of capsaicin and MTSET as indicated over current traces. Control TRPV1-C157A with 100 μM MTSET and co-application of 0.3 μM capsaicin. **(B)** Control TRPV1-C157A with 1 mM MTSET and co-application of 0.3 μM capsaicin. **(C)** TRPV1-C157A/M308C with 100 μM MTSET and co-application of 0.3 μM capsaicin. **(D)** TRPV1-C157A/M308C with 1 mM MTSET and co-application of 0.3 μM capsaicin.

Supplemental Figure 4

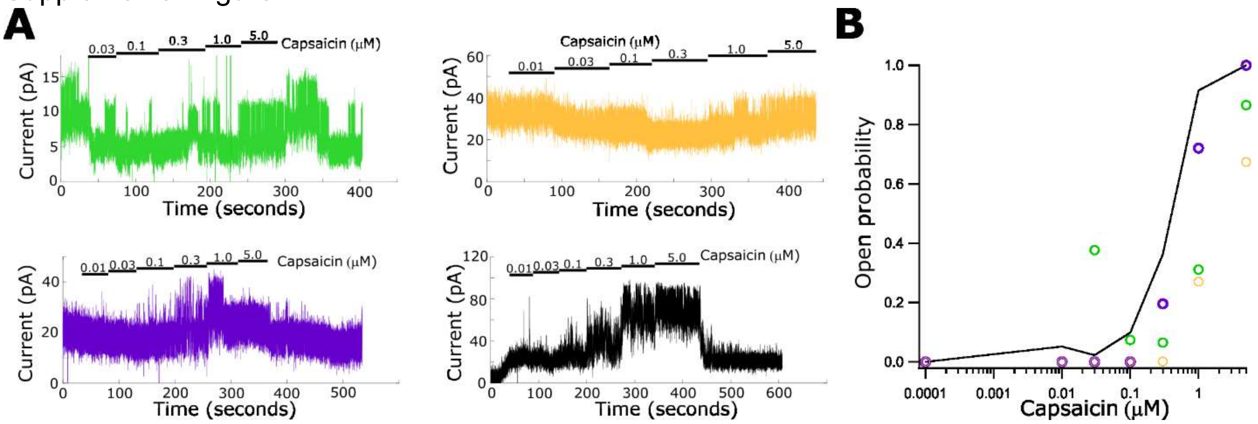


**Figure S4. TRPV1M308H dose response data with single channel and multi-channel patches. (A)** Currents collected in voltage sweeps with voltage set to +80mV. Because the voltage sweeps include a -80mV and +80mV segment, we used a macro to extract the +80mV segments and concatenate these in the presented and analyzed data (described in the Methods section). Current traces with single and double channel contributions are colored, and the multi-channel patch is black. Notable properties of the green trace is high activity for two channels in the wash buffer before and after capsaicin application. **(B)** Summary plot of the channel activity illustrated in data traces shown in (A). The multi-channel patch is plotted with a line and single channel data is plotted with colored markers. The purple trace shows two active channels until the application of the 5 μM solution when one channel becomes silent. The purple 5μM concentration data point in (B) is based on a single channel remaining active in the purple trace.

Supplementary Tables

Supplemental Table 1

| Uniprot ID | Entry name | Ensemble ID | Protein variant | SNV | AM Path. Score | Pathogenicity class |
|---|---|---|---|---|---|---|
| Q8NER1 | TRPV1_HUMAN | ENST00000399756.8 | p.Met309Ala | | 0.8691 | likely_pathogenic |
| Q8NER1 | TRPV1_HUMAN | ENST00000399756.8 | p.Met309Cys | | 0.9186 | likely_pathogenic |
| Q8NER1 | TRPV1_HUMAN | ENST00000399756.8 | p.Met309Asp | | 0.9988 | likely_pathogenic |
| Q8NER1 | TRPV1_HUMAN | ENST00000399756.8 | p.Met309Glu | | 0.9833 | likely_pathogenic |
| Q8NER1 | TRPV1_HUMAN | ENST00000399756.8 | p.Met309Phe | | 0.901 | likely_pathogenic |
| Q8NER1 | TRPV1_HUMAN | ENST00000399756.8 | p.Met309Gly | | 0.9855 | likely_pathogenic |
| Q8NER1 | TRPV1_HUMAN | ENST00000399756.8 | p.Met309His | | 0.9861 | likely_pathogenic |
| Q8NER1 | TRPV1_HUMAN | ENST00000399756.8 | p.Met309Ile | y | 0.7738 | likely_pathogenic |
| Q8NER1 | TRPV1_HUMAN | ENST00000399756.8 | p.Met309Lys | y | 0.9446 | likely_pathogenic |
| Q8NER1 | TRPV1_HUMAN | ENST00000399756.8 | p.Met309Leu | y | 0.5779 | likely_pathogenic |
| Q8NER1 | TRPV1_HUMAN | ENST00000399756.8 | p.Met309Asn | | 0.9753 | likely_pathogenic |
| Q8NER1 | TRPV1_HUMAN | ENST00000399756.8 | p.Met309Pro | | 0.9981 | likely_pathogenic |
| Q8NER1 | TRPV1_HUMAN | ENST00000399756.8 | p.Met309Gln | | 0.8607 | likely_pathogenic |
| Q8NER1 | TRPV1_HUMAN | ENST00000399756.8 | p.Met309Arg | y | 0.933 | likely_pathogenic |
| Q8NER1 | TRPV1_HUMAN | ENST00000399756.8 | p.Met309Ser | | 0.9081 | likely_pathogenic |
| Q8NER1 | TRPV1_HUMAN | ENST00000399756.8 | p.Met309Thr | y | 0.6428 | likely_pathogenic |
| Q8NER1 | TRPV1_HUMAN | ENST00000399756.8 | p.Met309Val | y | 0.1557 | likely_benign |
| Q8NER1 | TRPV1_HUMAN | ENST00000399756.8 | p.Met309Trp | | 0.9922 | likely_pathogenic |
| Q8NER1 | TRPV1_HUMAN | ENST00000399756.8 | p.Met309Tyr | | 0.9875 | likely_pathogenic |

Table S1. Excerpt of Alphamissense scores assigned to M309 in human TRPV1 (M308 in the rat TRPV1 sequence). SNV, indicates with “y” whether a single nucleotide variant is known to exist.
